# Enhancers regulate 3′ end processing activity to control expression of alternative 3′UTR isoforms

**DOI:** 10.1101/2020.08.17.254193

**Authors:** Buki Kwon, Mervin M. Fansler, Neil D. Patel, Jihye Lee, Weirui Ma, Christine Mayr

**Author notes:** Correspondence, Christine Mayr, Sloan Kettering Institute, 1275 York Ave, Box 303, New York, NY 10065. These authors contributed equally (Buki Kwon, Mervin M. Fansler, Neil D. Patel).

## Abstract

Multi-UTR genes are widely transcribed but show cell type-specific 3′UTR isoform expression. As transcriptional enhancers regulate mRNA expression, we investigated if they also regulate mRNA isoform expression. Deletion of an endogenous enhancer of a multi-UTR gene did not impair transcript production but prevented a switch in 3′UTR isoform expression. Also, the same enhancers are able to increase transcript production in the context of single-UTR gene promoters, but they increase 3′ end processing activity when paired with multi-UTR gene promoters. We show that transcription factors regulate processing activity of weak polyadenylation sites to control cell type-specific alternative 3′UTR isoform expression of widely expressed genes.

## Introduction

Most of our knowledge on gene expression regulation has been gained through analysis of genes that generate mRNAs with constitutive 3′UTRs, meaning that their pre-mRNAs are processed into mRNA isoforms with a single 3′UTR. This class of genes contains widely expressed housekeeping genes as well as the majority of developmentally regulated genes ^1-3^. Cell type-specific mRNA expression is known to be regulated by transcriptional enhancers ^1,2,4-6^. However, approximately half of human genes use alternative cleavage and polyadenylation (APA) to generate mRNA isoforms that encode the same protein but differ in their 3′UTR sequence ^3^. The vast majority of these genes are widely expressed, but they are characterized by tissue- and cell type-specific expression of specific 3′UTR isoforms. These genes are enriched in regulatory factors, including transcription factors, RNA-binding proteins, kinases, and ubiquitin enzymes ^3^. However, their mode of regulation is largely unknown, and it is currently unclear how cell type-specific expression of individual mRNA isoforms with unique 3′UTRs is achieved.

APA is developmentally regulated and can be dysregulated in disease ^7,8^. Inclusion of different regulatory elements in 3′UTRs influences mRNA stability, translation, and localization ^9,10^. A difference in 3′UTR sequence can also determine protein function as alternative 3′UTRs allow newly translated proteins to participate in alternative protein complexes ^11-15^. Alternative 3′UTR isoform usage is thought to be mostly regulated by differential expression of RNA-binding proteins, including polyadenylation and splicing factors, as their knock-down often changes 3′UTR isoform usage of hundreds of genes ^7,8,16-22^. However, genome-wide analyses of 3′UTR isoform expression across cell types and conditions revealed gene- and condition-specific changes in 3′UTR ratios, thus pointing to a more fine-grained regulation of APA ^3^.

According to the original definition, transcriptional enhancers are DNA sequences that increase the expression of a reporter gene ^4-6,23^. Currently, increased gene expression is often used interchangeably with increased transcription, thus implying that enhancers mostly affect transcript production ^6,24^. However, the generation of mature mRNAs requires pre-mRNA production and processing which includes splicing and 3′ end cleavage and polyadenylation (CPA) ^8,25^. Therefore, when disregarding the contribution of mRNA stability, mRNA production of unspliced transcripts is largely determined by the number of produced transcripts and by the 3′ end processing activity that we call here CPA activity.

Under physiological conditions it is currently difficult to disentangle the contribution of transcript production and transcript processing to the expression level of single-UTR genes. However, viral infection or osmotic stress impair transcript processing of cellular genes and lead to massive read-through transcription downstream of polyadenylation signals (PAS), thus illustrating the crucial contribution of transcript processing ^26-29^. Moreover, point mutations or genetic variants that occur in PAS or in their surrounding sequence elements reveal the contribution of 3′ end processing activity to mRNA expression ^30-36^. Such mutations result in 1.5 to 2-fold differences in steady-state mRNA levels which is sufficient to cause disease phenotypes, including thalassemia, thrombophilia, or cancer predisposition ^30-32,34,35^.

Here, we set out to investigate if cell type- or condition-specific expression of 3′UTR isoforms is regulated by transcriptional enhancers. We found that deletion of an endogenous enhancer associated with a multi-UTR gene did not reduce transcript production but impaired CPA activity at a proximal and weak polyadenylation site. The enhancer-dependent processing activity regulation was mediated by transcription factors, including MYC and NF-κB that are known to bind to the enhancer. Enhancer-mediated regulation of 3′UTR isoform expression is widespread as endogenous, cell type-specific enhancers significantly associate with genes that exclusively upregulate their 3′UTR isoform expression in a cell type-specific manner. Our data indicate that transcriptional enhancers regulate both aspects of mature mRNA generation, namely transcript production and 3′ end processing to regulate mRNA and mRNA isoform expression in a cell type- and condition-specific manner.

## Results

### The *PTEN* enhancer induces a switch in 3′UTR isoform expression of endogenous *PTEN*

*PTEN* is a tumor-suppressor gene whose expression is altered in a large fraction of cancers. Cells are very sensitive to *PTEN* dosage as even a small decrease in PTEN expression is cancer-promoting ^37^. The *PTEN* gene generates multiple mRNA isoforms with alternative 3′UTRs that encode the same protein. In our previous 3′UTR isoform expression study, *PTEN* was among the top genes with extensive differences in alternative 3′UTR isoform usage across cell lines and tissues ^3^. To obtain a better understanding of *PTEN* expression regulation, we applied CRISPR technology to delete the promoter-proximal *PTEN* enhancer in the breast cancer cell line MCF7 which expresses wild-type *PTEN* ^38^. The boundaries of the *PTEN* enhancer were determined using ChIP-seq data on transcription factor binding sites and acetylated H3K27 levels (Fig. 1a) ^39-42^. We deleted the *PTEN* enhancer with a pair of guide RNAs and obtained two control clones with the wild-type (WT) enhancer sequence and two ‘delta enhancer’ (dE) clones with a heterozygous deletion in the region of the *PTEN* enhancer (Fig. 1b and Extended Data Fig. 1a,b).

**Fig. 1:**
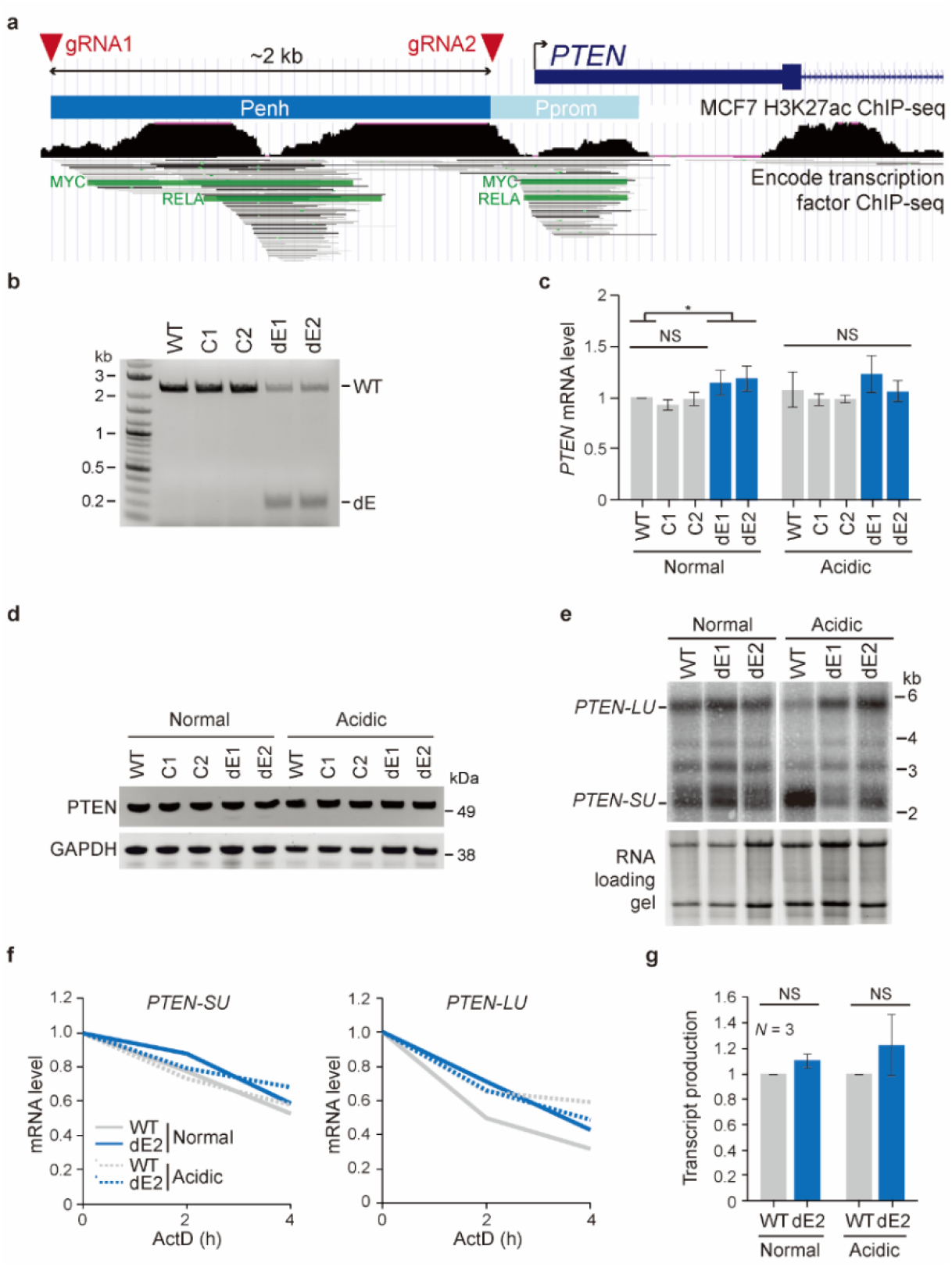
The *PTEN* enhancer regulates alternative 3′UTR isoform expression of *PTEN*. **(a)** UCSC genome browser snapshot showing the *PTEN* genomic locus around the transcriptional start site (arrow). The *PTEN* enhancer (Penh) was deleted using the indicated guide RNAs (red arrow heads). Among transcription factor binding sites identified by ChIP-seq, MYC and RELA binding sites are highlighted. *PTEN* promoter, Pprom. **(b)** Genotyping PCR was performed using a primer pair flanking the *PTEN* enhancer. Shown is a full-length product (WT) in parental MCF7 cells and wild-type clones (C1, C2) and an additional shorter product indicating heterozygous deletion of the enhancer in dE1 and dE2 clones. **(c)** *PTEN* mRNA expression measured by RT-qPCR in the indicated samples. Data are shown as mean ± std. of *N* = 4 biological replicates for C1 and C2 and *N* = 8 biological replicates for WT, dE1, and dE2 after normalization to *RPL19*. T-test for independent samples was performed. *, *P* < 0.02; NS, not significant. **(d)** Representative western blot showing steady-state PTEN protein level in the indicated samples. GAPDH serves as loading control. **(e)** Representative northern blot showing *PTEN* mRNA isoforms in the indicated samples. The RNA gel is shown as loading control. *SU*, short 3′UTR; *LU*, long 3′UTR. **(f)** Quantification of *PTEN* mRNA isoform expression at two time points after inhibition of transcription with actinomycin D (ActD) in the indicated samples. The values were obtained by northern blot analysis (shown in Extended Data Fig. 1f) and were normalized to the zero-hour time points. **(g)** Metabolic labeling with 4-thiouridine (4sU) was used to enrich newly transcribed mRNAs. The newly transcribed RNAs were thiol-alkylated and biotinylated, followed by Streptavidin pull-down. The fraction of newly transcribed over total *PTEN* transcripts is shown for the indicated samples and was measured using RT-qPCR with a primer pair in the first intron. T-test for independent samples was performed. NS, not significant.

Heterozygous deletion of the enhancer increased steady-state *PTEN* mRNA level by only 1.17-fold and did not affect protein level (Fig. 1c,d). We hypothesized that enhancer activation may be necessary to observe an effect. The *PTEN* enhancer contains conserved MYC binding sites and canonical NF-κB binding sites (Fig. 1a and Extended Data Fig. 1c,d) ^43,44^. As cytoplasmic acidification in MCF7 cells was previously shown to increase NF-κB and MYC activity ^45,46^, we cultivated the cells in normal or acidified media (pH = 6.5), but did not observe an enhancer-mediated difference in *PTEN* mRNA and protein levels (Fig. 1c,d).

Promoters were previously implicated in the regulation of mRNA processing ^47-56^. Therefore, we investigated if deletion of the enhancer would change alternative 3′UTR isoform expression of *PTEN*. Enhancer deletion had little effect on 3′UTR isoform expression under normal cultivation conditions (Fig. 1e and Extended Data Fig. 1e). However, in acidified conditions, we observed a striking switch in 3′UTR isoform expression with increased expression of the short 3′UTR (*SU*) isoform of *PTEN* which was fully abrogated in cells lacking the enhancer (Fig. 1e and Extended Data Fig. 1e). This result demonstrates that the *PTEN* enhancer is required for a 3′UTR isoform change of *PTEN*. The switch in 3′UTR isoform expression is either caused by a change in alternative PAS usage or by preferential degradation of the long 3′UTR (*LU*) isoform. However, degradation of the *LU* isoform would need to be accompanied by increased transcript production to account for the upregulated *SU* expression.

To identify the mechanism by which the enhancer controls the switch in 3′UTR isoform expression, we measured transcript production and stability. We inhibited transcription with actinomycin D and performed northern blot analysis at different time points to measure stability of the alternative 3′UTR isoforms (Fig. 1f and Extended Data Fig. 1f). We did not detect an enhancer- or condition-specific difference in stability of the mRNA isoforms (Fig. 1f and Extended Data Fig. 1f). We then used metabolic labeling with 4-thiouridine to examine an enhancer-dependent change in transcript production before and after media acidification. We did not observe a significant difference in *PTEN* pre-mRNA production between WT and dE mutant cells or between normal and acidified conditions (Fig. 1g). Enhancers are also known to regulate alternative splicing ^55^, but we did not observe enhancer-dependent alternative splicing of *PTEN* (Fig. 1e and Extended Data Fig. 1e,g). As the *PTEN* enhancer was required for a switch in 3′UTR isoform expression without substantially regulating transcript production or causing differential stability of the alternative mRNA transcripts, our data suggest that it regulates CPA activity.

### The *PTEN* enhancer increases 3′ end processing activity of weak PAS in a reporter system

Processing activity of endogenous transcripts cannot be fully disentangled from transcript production and stability. To investigate if transcriptional enhancers indeed control CPA activity, we developed a reporter system. Our luciferase reporter system allows us to separately investigate enhancer-dependent transcript production and transcript processing. The polyadenylation signal (PAS) derived from the SV40 late transcript is one of the strongest known PAS ^57,58^. When used for termination of a luciferase reporter construct it results in processing of all produced transcripts (Fig. 2a) ^57,58^. Therefore, the SV40 PAS reporter construct measures transcriptional activity of the promoter, a system that has been widely used to measure transcriptional activity ^59^. To assess enhancer-dependent CPA activity, we measured luciferase activity of a reporter construct that is nearly identical except that it is terminated by the proximal PAS (PPAS) of *PTEN* instead of the SV40 PAS. As only processed transcripts contribute to luciferase activity, the ratio of luciferase activities obtained from the PTEN PPAS reporter over the SV40 PAS reporter represents the relative CPA activity of the PTEN PPAS when driven by a specific promoter (Fig. 2a). In this reporter system, CPA activity corresponds to luciferase activity if the different polyadenylation sites do not influence stability of the reporters. To minimize the elements that may affect mRNA stability we used minimal polyadenylation sites that only contain ∼100 base pairs of surrounding sequence to ensure proper 3′ end processing (Supplementary Table 1a) ^25^.

**Fig. 2:**
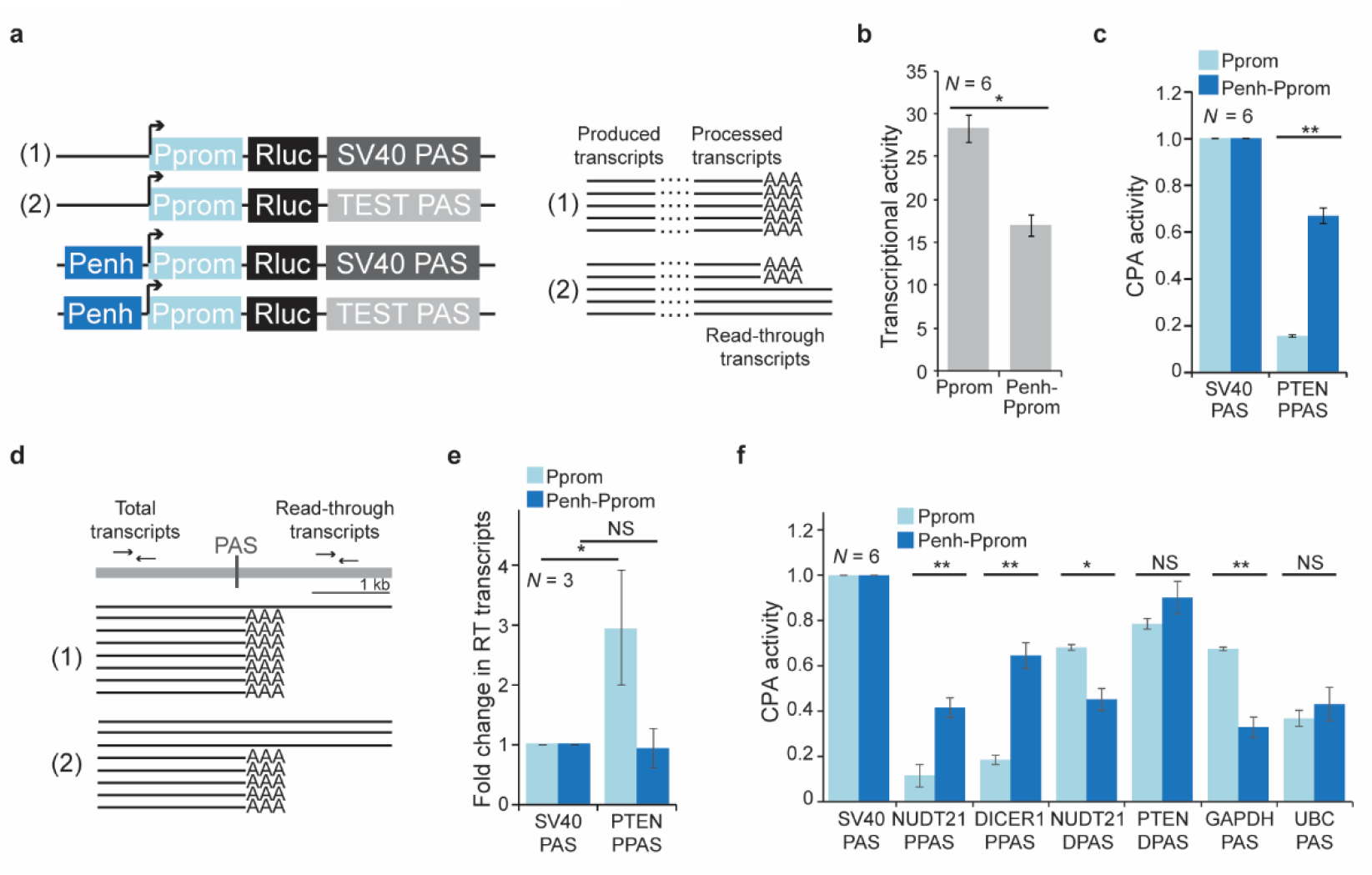
The *PTEN* enhancer increases CPA activity of weak polyadenylation sites. **(a)** Schematic of luciferase reporter constructs to investigate enhancer-dependent transcriptional activity and CPA activity. The transcription start site is indicated by the arrow. Rluc, Renilla luciferase. Expected reporter transcript production and processing for the indicated constructs. In reporter (1), the strong SV40 PAS cleaves all produced transcripts (indicated by AAA to denote a poly(A) tail) and measures transcriptional activity. In reporter (2) a weaker PAS does not cleave all produced transcripts, thus resulting in read-through transcripts. The relative CPA activity of a test PAS corresponds to the ratio of the luciferase activities obtained from the test PAS reporter over the SV40 PAS reporter when transcribed from the same promoter. **(b)** Transcriptional activity of the Pprom reporter in the presence (Penh-Pprom) or absence (Pprom) of the *PTEN* enhancer obtained by luciferase activity of the SV40 PAS reporters. Transcriptional activity represents renilla luciferase activity that was normalized by firefly luciferase activity. Shown is mean ± std. T-test for independent samples was performed; *, *P* = 0.002. **(c)** Relative CPA activity of the PPAS of PTEN when transcribed from the *PTEN* promoter in the absence or presence of the *PTEN* enhancer. Shown is mean ± std. T-test for independent samples was performed; **, *P* = 1 × 10^−8^; NS, not significant. **(d)** Schematic for measuring read-through transcription of the reporter constructs shown in (a). A primer pair located upstream of the PAS measures the total number of transcripts produced, whereas a primer pair located downstream of the PAS measures the number of read-through transcripts. **(e)** Fold change in read-through (RT) transcripts obtained from the indicated reporter constructs. Shown is mean ± std. T-test for independent samples was performed; *, *P* = 0.04; NS, not significant. **(f)** As in (c), but additional PAS are shown. DPAS, distal PAS. Shown is mean ± std. T-test for independent samples was performed; **, *P* = 1 × 10^−6^; *, *P* = 0.001. NS, not significant.

We measured transcriptional activity of the *PTEN* promoter (Pprom) and observed a slight decrease in transcriptional activity (1.6-fold) in the presence of the enhancer (Penh-Pprom; Fig. 2b). The decrease may be due to the high transcriptional activity of the isolated *PTEN* promoter and the fact that some transcription factors act as repressors. However, typical enhancers increase transcriptional activity ^6,23^. When we analyzed the effect of the *PTEN* enhancer in the context of two weaker core promoters, it indeed increased transcriptional activity, indicating that it acts as a transcriptional enhancer following the original definition (Extended Data Fig. 2a,b) ^6,23^. Intriguingly, the addition of the enhancer increased luciferase activity of the PTEN PPAS reporter 4-fold (Fig. 2c). To investigate if the increase in luciferase activity is the result of an enhancer-dependent increase in 3′ end processing activity, we measured transcript abundance upstream and downstream of the PAS which enables detection of potential differences in the levels of read-through transcripts in the four reporter constructs (Fig. 2d). All constructs showed a comparable amount of read-through transcription with exception of the PTEN PPAS reporter when transcribed from the *PTEN* promoter lacking the enhancer (Fig. 2e). We observed 3-fold more read-through in this reporter which is consistent with its lower luciferase activity (Fig. 2c) and supports enhancer-dependent regulation of CPA activity of this weak polyadenylation site.

We performed additional control experiments to gain a better understanding of enhancer-dependent reporter regulation. We did not observe enhancer- or PAS-dependent differences in mRNA stability of the reporters, indicating that the difference in luciferase activity correlates with CPA activity (Extended Data Fig. 2c). The addition of the enhancer did not change the transcription start site of the reporter (Extended Data Fig. 2d), indicating that the mature mRNAs produced from the *PTEN* promoter in the presence or absence of the *PTEN* enhancer are identical. Enhancers regulate transcription independently of their orientation and can be located up- or downstream of genes ^6,42^. When we placed the reverse complement of the *PTEN* enhancer downstream of the PAS it enhanced transcription and enhanced PAS cleavage (Extended Data Fig. 2e-g). Taken together, these data suggest that regulation of 3′ end processing activity is a *bona fide* activity of transcriptional enhancers.

Next, we assessed if the enhancer controls processing activity of additional polyadenylation sites (Supplementary Table 1a). We tested two PPAS (derived from *NUDT21* and *DICER1*), two distal PAS (DPAS; derived from *PTEN* and *NUDT21*), and two PAS derived from housekeeping genes (*GAPDH* and *UBC*) that generate constitutive 3′UTRs ^3^. The CPA activity of all proximal PAS was weaker than that of other PAS in the absence of the enhancer (Fig. 2f). However, the enhancer increased PPAS cleavage activity by up to 3.6-fold (Fig. 2f). In contrast, the enhancer did not change cleavage activity of stronger, non-proximal polyadenylation sites in a coordinated manner (Fig. 2f). These observations suggest that in the absence of an enhancer CPA activity largely depends on the intrinsic strength of a PAS ^60^, but intrinsically weak polyadenylation sites can have high *in vivo* cleavage activity when transcribed from promoters with active enhancers.

### A distal enhancer also regulates CPA activity of weak polyadenylation sites

We then asked if other enhancers are also capable of regulating CPA activity and set out to test the enhancer of the *NUDT21* gene. The *NUDT21* gene encodes an important polyadenylation factor that changes CPA of hundreds of genes, when knocked-down ^18,20^. Moreover, it generates alternative 3′UTRs and similar to *PTEN*, its 3′UTR isoforms are extensively regulated across samples ^3^. We searched for ChIP-seq peaks with high H3K27 acetylation level in the vicinity of the *NUDT21* gene, as high H3K27 acetylation levels are characteristic for enhancers ^6,42^. H3K27 acetylation levels in the promoter-proximal region of *NUDT21* were only intermediate, but we detected a region with very high acetylation levels 80 kb downstream of the *NUDT21* gene. We cloned 2 kb of this region and called it distal enhancer (Denh; Fig. 3a) as we currently have no evidence that this region is an enhancer of the *NUDT21* gene. To test if the distal enhancer is functional, we measured enhancer-dependent transcriptional activity of the *GAPDH* promoter (Gprom) which drives expression of a single-UTR gene (Fig. 3b) ^3^. The distal enhancer upregulated transcriptional activity by more than 4-fold, thus acting as a transcriptional enhancer (Fig. 3c). However, the distal enhancer had no influence on CPA activity in the context of the Gprom (Fig. 3d). Similar results were obtained when using the *PTEN* enhancer in the context of the Gprom (Extended Data Fig. 2b,h). In contrast, in the context of two promoters derived from multi-UTR genes, such as the *PTEN* or *NUDT21* promoters (Nprom), the distal enhancer did not significantly affect transcriptional activity (Fig. 3b,e), but instead increased PTEN PPAS cleavage activity between 3.4 and 5.3-fold (Fig. 3f,g). As the distal enhancer did not affect processing activity of a stronger PAS, our results indicate that transcriptional enhancers regulate CPA activity of weak polyadenylation sites in the context of multi-UTR promoters (Fig. 3f,g).

**Fig. 3:**
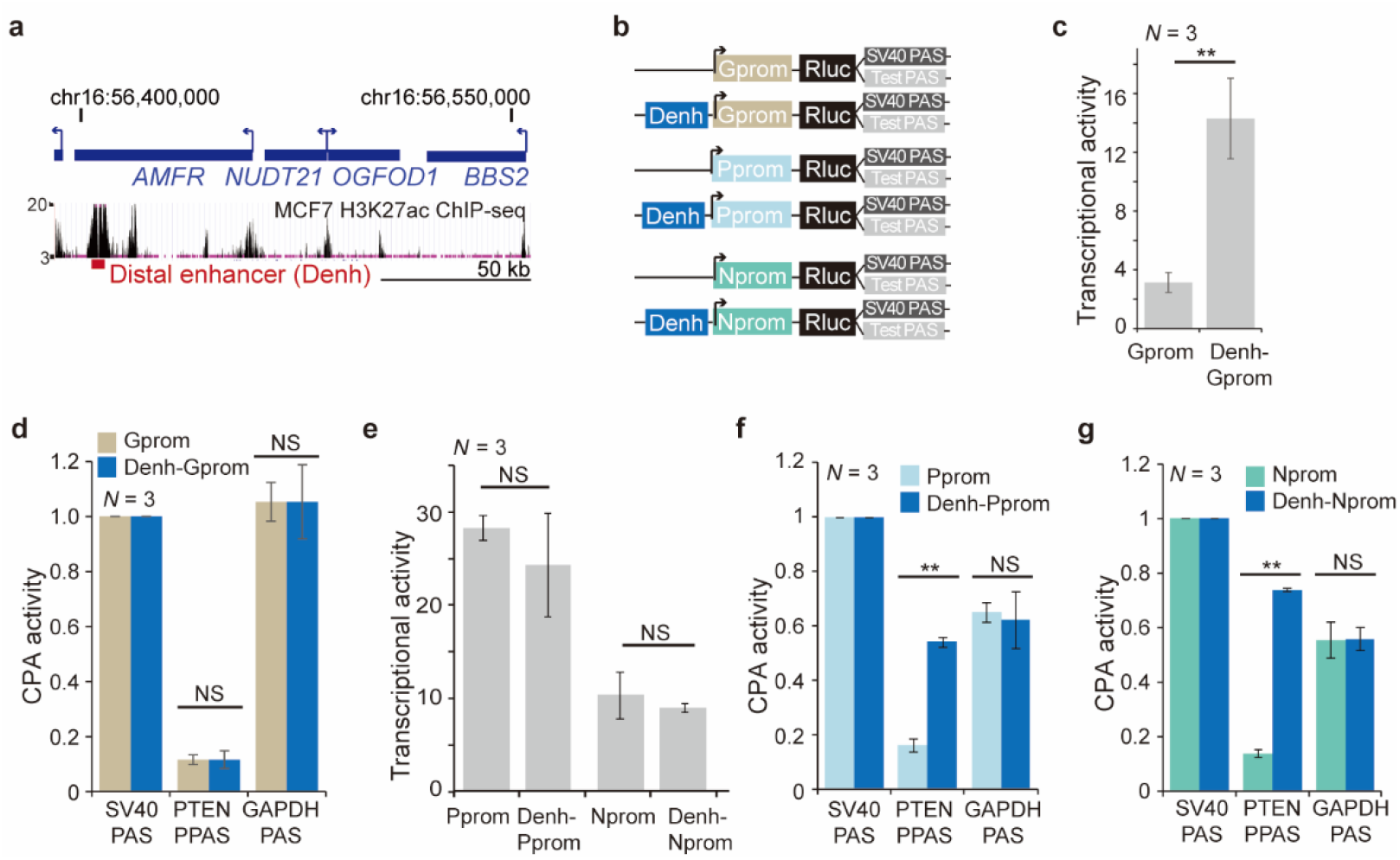
A distal enhancer regulates CPA activity of weak polyadenylation sites. **(a)** UCSC genome browser snapshot showing the genomic context of the *NUDT21* gene locus. The region with the local maximum of acetylated H3K27 measured by ChIP-seq in MCF7 cells was defined as distal enhancer (Denh). See Supplementary Table 1 for coordinates. **(b)** Schematic of reporter constructs used to investigate enhancer-dependent CPA activity in the context of three promoters. The *GAPDH* promoter (Gprom) drives a single-UTR gene, whereas the Pprom and *NUDT21* (Nprom) promoters drive multi-UTR genes. Shown as in Fig. 2a. **(c)** Enhancer-dependent transcriptional activity of the Gprom. As in Fig. 2b. **, *P* = 0.0005. **(d)** Enhancer-dependent CPA activity in the context of the Gprom. As in Fig. 2c. NS, not significant. **(e)** As in (c), but enhancer-dependent transcriptional activity of two multi-UTR gene promoters is shown. **(f)** As in (d), but enhancer-dependent CPA activity in the context of the Pprom is shown. **, *P* = 1 × 10^−8^. **(g)** As in (d), but enhancer-dependent CPA activity in the context of the Nprom is shown. **, *P* = 1 × 10^−5^.

### Transcription factors can regulate CPA activity without affecting transcriptional activity

To better understand how enhancers regulate CPA activity in the context of multi-UTR promoters, we set out to identify the transcription factors that mediate enhancer-dependent regulation of 3′ end processing activity in the context of the *PTEN* promoter. We performed a small-scale shRNA screen by knocking-down (KD) individual transcription factors that are expressed in MCF7 cells and that were shown by ChIP-seq to bind to the *PTEN* enhancer (Fig. 4a, Extended Data Fig. 3 and Supplementary Table 2) ^40^. We measured transcriptional and CPA activities in the context of the Penh-Pprom in control KD and transcription factor KD samples (Fig. 4b). As positive control, we knocked-down the polyadenylation factor FIP1L1, which was shown previously to be required for PPAS usage ^17^. KD of FIP1L1 decreased PTEN PPAS usage from 0.6 to 0.36 without affecting transcriptional activity (Fig. 4c black and Extended Data Fig. 4a,b). Sixteen out of 21 tested transcription factors significantly changed CPA activity. Whereas a few of them simultaneously changed transcriptional and CPA activities (Fig. 4c purple), we obtained the striking result that KD of ten individual transcription factors only regulated CPA activity without exerting a strong (less than 1.7-fold) effect on transcriptional activity of the reporter (Fig. 4c red). These transcription factors included RELA (NF-κB p65) and MYC (Fig. 4c, Table 1 and Supplementary Table 2).

**Table 1.** Transcription factors and co-activators that regulate CPA activity of weak PAS.

| Activity | Transcription factors | Co-activators |
| --- | --- | --- |
| Regulation of CPA activity only | FOS, FOXA1, GABPA, IRF1, JUN, JUND, MYC, NFYB, RELA, TCF12 | ING3, SMC3 |
| Regulation of CPA and transcription | EGR1, ELF1, RFX5, RXRA, TFAP2A, YY1 | ACTL6A, BARD1, BRCA1, BRD8, CDC73, CDK9, EP300, EP400, GTF2F1, KAT2B, KAT5, MED1, MORF4L1, NELFB, NELFE, NPM1, SIN3A, SMARCC1, SSRP1, SUPT16H, SUPT4H1, SUPT5H, TAF1, TBP, TCERG1, USP22 |

**Fig. 4:**
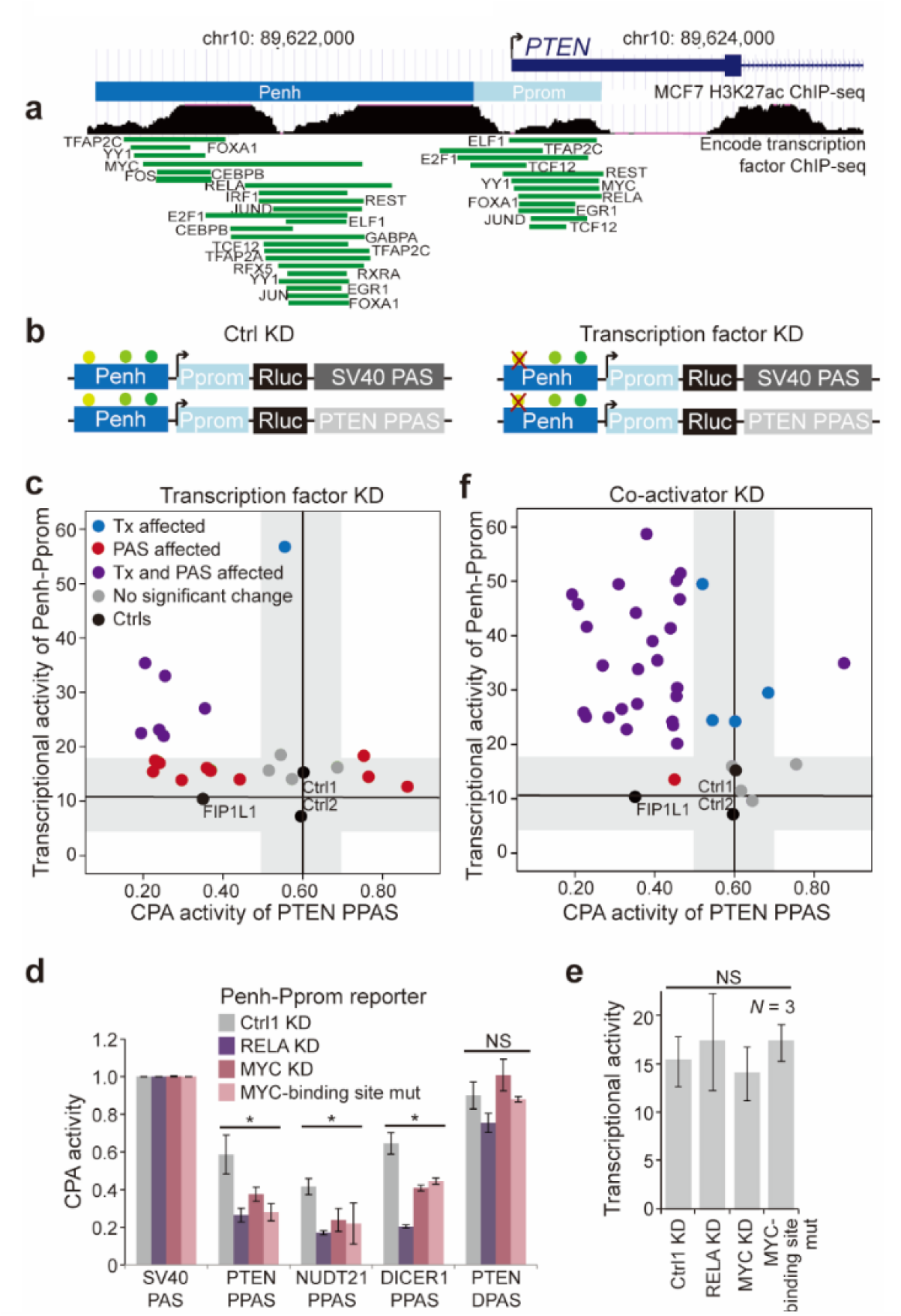
Transcription factors regulate CPA activity without affecting transcriptional activity. **(a)** As in Fig. 1a but shown are the binding sites of transcription factors that were knocked down individually. **(b)** Schematic of reporter constructs to identify transcription factors that regulate CPA activity in the context of the Penh-Pprom reporter. Ctrl KD, control knock-down. **(c)** Summary of transcriptional activity and PTEN PPAS CPA activity obtained in the shRNA screen. Shown is the mean in ctrl KD and transcription factor KD samples for the Penh-Pprom reporter. The horizontal/vertical grey areas denote no significant change in transcriptional and CPA activities, respectively. Tx, transcription. See Extended Data Fig. 4a,b and Supplementary Table 2 for complete data. **(d)** Mean ± std of CPA activity of additional PAS when transcribed from the Penh-Pprom reporter. Shown is the effect of transcription factor KD or mutation of MYC-binding sites (MYC binding site mut) in the Penh. T-test for independent samples was performed; *, *P* < 0.03, NS, not significant. **(e)** Transcriptional activity of the Penh-Pprom reporter upon transcription factor KD or after mutation of MYC-binding sites is shown as in Fig. 2b. **(f)** As in (c), but KD of co-activators is shown.

### Mutation of MYC binding sites in the enhancer decreases CPA activity

RELA and MYC KD also decreased CPA activity of additional PPAS but did not significantly affect cleavage activity of the strong PTEN DPAS (Fig. 4d and Extended Data Fig. 4c). Similar results were obtained after mutation of the two highly conserved MYC-binding sites in the *PTEN* enhancer (Extended Data Fig. 1c,d). Mutation of the MYC-binding sites had no influence on transcriptional activity (Fig. 4e), did not affect CPA activity of the strong DPAS of PTEN, but decreased CPA activity of weak PPAS, thus phenocopying the effect of MYC KD (Fig. 4d). These results suggest that binding of transcription factors to conserved motifs in the *PTEN* enhancer regulates CPA activity of weak proximal PAS.

### Co-activators simultaneously regulate transcriptional and PAS cleavage activity

Binding of transcription factors to enhancers results in recruitment of co-activators to promoters ^61-63^. Co-activators include components of the Mediator complex, the general transcription machinery, transcription elongation factors, and histone acetyltransferases ^61-63^. In contrast to transcription factors, silencing of the majority of co-activators (26/35) changed transcriptional activity and CPA activity at the same time (Fig. 4f, purple, Table 1, Extended Data Fig. 4a,b and Supplementary Table 2). Moreover, impaired enhancer activation upon KD of histone acetyl transferases such as TIP60 and PCAF (encoded by *KAT5* and *KAT2B*, respectively) reduced CPA activity of several weak proximal polyadenylation sites in the context of the *PTEN* enhancer (Extended Data Fig. 4d). KD of histone acetyltransferases also decreased PTEN PPAS CPA activity in the context of the distal enhancer but had no effect on CPA activity in reporters that lack the enhancer (Extended Data Fig. 4e-h). These results suggest that regulation of cleavage activity of weak polyadenylation sites by active enhancers has the potential to be widespread as two out of two tested enhancers regulated 3′ end processing activity in the context of the reporter.

### Cell type-specific enhancers preferentially associate with genes that upregulate *SU* isoforms in a cell type-specific manner

Next, we set out to investigate if endogenous enhancers are widespread regulators of CPA activity. The analysis of multi-UTR genes allows us to distinguish transcriptional and CPA activity. Transcriptional upregulation will increase *SU* and *LU* isoform expression to a similar extent, whereas increased CPA activity of proximal polyadenylation sites will only increase *SU*, but not *LU* isoform expression.

To examine if cell type-specific enhancers are associated with changes in 3′UTR isoform expression, we used a dataset that mapped erythroblast-specific enhancers and associated them with individual genes ^64^. To identify genes that become upregulated in erythroblasts, we compared gene expression between erythroblasts and hematopoietic stem cells ^65,66^. We analyzed single- and multi-UTR genes separately, but as expected, we observed that genes that increase their expression in erythroblasts preferentially associate with enhancers that are active in erythroblasts (Fig. 5a,b) ^67^. The association with erythroblast-specific enhancers was not observed for genes whose expression did not increase (Fig. 5a,b).

**Fig. 5:**
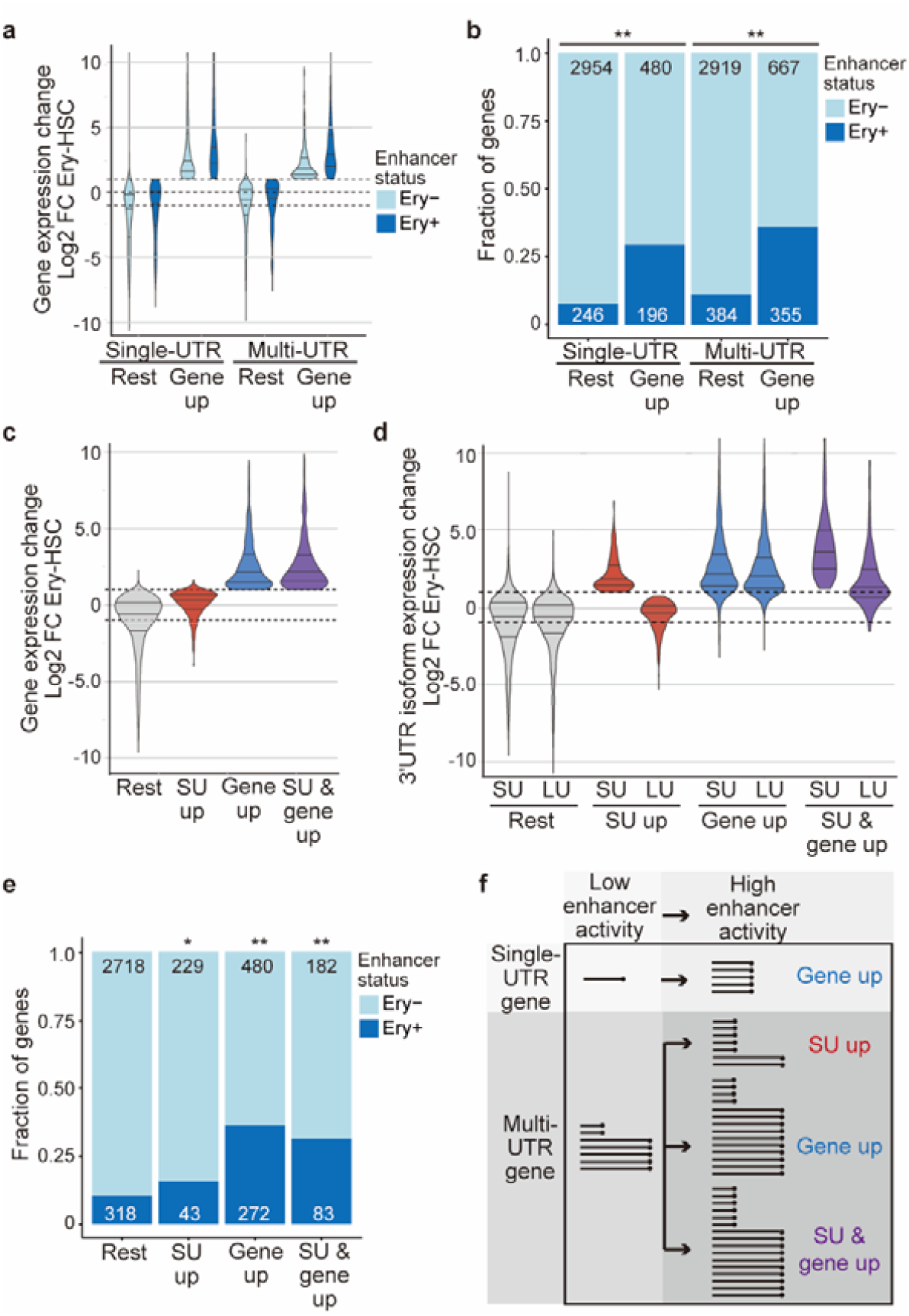
Cell type-specific enhancers are associated with genes that upregulate *SU* isoforms. **(a)** Fold change (FC) in gene expression between erythroblasts (Ery) and hematopoietic stem cells (HSC) is shown for single- and multi-UTR genes separately. The genes that associate with erythroblast-specific enhancers (Ery+) are shown separately. Violin plots denote median, 25^th^ and 75^th^ percentiles. The number of genes in each group is shown in (b). **(b)** Fraction of genes associated with erythroblast-specific enhancers (Ery+) in the groups from (a). Chi-square tests were performed. Single-UTR genes: X^2^ = 249, **, *P* < 10^−16^. Multi-UTR genes: X^2^ = 293, **, *P* < 10^−16^. Ery-, genes without an erythroblast-specific enhancer in the vicinity. **(c)** FC in gene expression in the indicated groups derived from multi-UTR genes is shown as in (a). The number of genes in each group is shown in (e). **(d)** FC in 3′UTR isoform expression shown for *SU* and *LU* isoforms in the groups from (c). **(e)** Fraction of genes associated erythroblast-specific enhancers (Ery+) in the groups from (c). Chi-square tests were performed. SU up, X^2^ = 7.3, *, *P* = 0.0069; gene up, X^2^ = 303, **, *P* < 10^−16^; both up, X^2^ = 99, **, *P* < 10^−16^. **(f)** Model showing enhancer-mediated increase in gene expression for single-UTR genes (top). Enhancer-mediated effect for multi-UTR genes can result in increased 3′UTR isoform expression, increased gene expression or an increase in both parameters. The lines with the black dots signify processed transcripts that contain a poly(A) tail.

Next, we focused on all multi-UTR genes and separated them into four groups. (1) *SU* isoform expression increases significantly, but overall gene expression does not (‘SU up’), (2) gene expression increases significantly with both *SU* and *LU* isoforms increase in a similar manner (‘gene up’), (3) *SU* isoform and gene expression increase significantly (‘*SU* and gene up’), and (4) the rest of the multi-UTR genes (Fig. 5c,d). As expected, both groups whose genes increased their gene expression were significantly associated with cell type-specific enhancers (Fig. 5e). Importantly, also genes with exclusive upregulation of their *SU* isoforms without leading to a significant upregulation in overall gene expression were significantly associated with cell type-specific enhancers (Fig. 5e). Taken together, our analysis revealed that cell type-specific enhancers are frequently associated with 3′UTR isoform changes as 32% of enhancers that were linked to expression changes were associated with significant 3′UTR isoform changes.

### Model of enhancer-mediated control of 3′UTR isoform expression

It is well-known that active enhancers upregulate gene expression of single-UTR genes in a cell type-specific manner (Fig. 5f, top) ^67^. In contrast, multi-UTR genes are usually transcribed in the majority of cell types ^3^. We found that enhancer activation of multi-UTR genes can have three potential outcomes. It can result in upregulation of gene expression with *SU* and *LU* isoforms increasing similarly. It can result in a combination of change in gene and isoform expression or it can result in upregulation of *SU* isoforms without a change in gene expression (Fig. 5f, bottom).

## Discussion

Mature mRNA production is determined by the extent of transcript production and transcript processing. Here, we show that transcriptional enhancers regulate both stages of mature mRNA production, but they differentially control them for different classes of genes. At single-UTR genes, enhancers increase transcript production, whereas at multi-UTR genes, transcriptional enhancers increase transcript production or transcript processing, thus either resulting in a gene expression change or in a cell type- or condition-specific change in 3′UTR isoform expression.

With our newly developed reporter assay, we were able to measure separately the two parameters of mRNA production and our results are consistent with the expression pattern observed for endogenous single- or multi-UTR genes. Whereas single-UTR genes are often transcribed in a cell type-specific manner, meaning that their expression is ‘off’ in some cell types and ‘on’ in others, most multi-UTR genes are always ‘on’ as they are transcribed in the majority of cell types ^3^. Despite their near ubiquitous expression, they encode regulatory factors and show cell type-specific 3′UTR isoform expression ^3^. This expression pattern was mirrored in the reporter assay, where two transcriptional enhancers increase transcript production when the reporter gene is driven by a single-UTR gene promoter. In contrast, the same enhancers did not affect overall transcription but instead increased the expression of a transcript terminated by a weak polyadenylation site when the reporter gene was driven by promoters derived from multi-UTR genes (Fig. 3). As the enhancer-dependent increase in isoform expression was associated with decreased read-through transcription downstream of weak polyadenylation sites, we conclude that enhancers regulate 3′ end processing activity (Fig. 2). Our reporter data are further supported by results obtained at endogenous gene loci, where enhancer deletion of a multi-UTR gene did not change gene expression but altered 3′UTR isoform expression (Fig. 1). Moreover, we found that cell type-specific enhancers were significantly associated with genes that exclusively upregulated their short 3′UTR isoforms without leading to an overall increase in gene expression (Fig. 5).

Our study further revealed that transcription factors and co-activators are responsible for increased transcript processing at weak polyadenylation sites and subsequent upregulation of mRNA isoform expression (Fig. 4). However, the exact mechanism by which enhancers regulate CPA activity are currently unknown. Based on the literature, several potential mechanisms exist that are not mutually exclusive. It is established that RNA-binding proteins that bind to the polyadenylation signal and the surrounding sequence determine 3′ end cleavage and polyadenylation ^7,8,16-22,56,58,68^. One model by which these RNA-binding proteins bind to polyadenylation sites is through the promoter loading model: Active enhancers recruit these factors to promoters which allows them to travel with RNA polymerase II and to bind to a newly transcribed polyadenylation site, thus increasing 3′ end processing activity locally ^47-55^. Such a mechanism has been proposed for promoter-dependent regulation of post-transcriptional processes in yeast, including the regulation of mRNA stability, cytoplasmic localization, and translation ^69-73^. This model is supported by the presence of a variety of RNA-binding proteins at 80% of human promoters and by the observation that many RNA-binding proteins bind to transcription factors ^52,53^.

Another model suggests that active enhancers regulate transcription elongation rate which could result in differential usage of polyadenylation sites ^74-76^. This is supported by our data showing that silencing of several factors involved in transcription elongation, including SPT4/5, NELF, and subunits of the PAF complex affect CPA activity of weak polyadenylation sites (Table 1). Finally, the integrator complex has been shown to associate with active enhancers and increases enhancer-promoter communication ^77^. Decreased expression of INTS11, the catalytic subunit of the complex, promotes read-through transcription at polyadenylation sites and shifts alternative isoform expression towards the distal isoform ^29,78^. This suggests that increased association of integrator at active enhancers could prevent read-through and could increase CPA activity. Integrator is also known to regulate transcription elongation ^79,80^, but it is currently unclear if its role in transcription elongation is required for integrator-dependent regulation of 3′ end processing of protein-coding genes.

What is the functional relevance of enhancer-mediated regulation of CPA activity? A recent large-scale study applied CRISPRi to identify functional enhancers and found that only 10% of tested enhancers showed any evidence of enhancer-mediated regulation of mRNA expression levels ^81^. Since in our study 11% of functional enhancers were associated with increased *SU* isoform expression, but not with increased gene expression, we suggest future studies include 3′UTR isoform expression as additional read-out for the testing of functional transcriptional enhancers.

Although APA is widespread and is regulated in a cell type- and condition-specific manner, for most genes the consequences of a change in alternative 3′UTR isoform expression are unknown ^3,7,8^. Nevertheless, striking examples exist in the literature that revealed that altering alternative 3′UTR isoform expression can result in substantial changes in protein expression caused by 3′UTR-dependent control of mRNA stability or translation ^33,82,83^. However, alterations in alternative 3′UTR isoform expression often do not change overall protein levels (Fig. 1) ^12,14,20,84,85^. For those cases, it has been demonstrated that isoform-specific differences in protein localization or function occur through 3′UTR-mediated formation of alternative protein complexes ^11-15^. For example, the ubiquitin ligase BIRC3 switches from predominant *SU* isoform expression in normal B cells to *LU* isoform expression in malignant B cells. Normal B cells mostly form protein complexes that are independent of *LU* isoforms and they mediate BIRC3’s tumor-suppressive functions. In contrast, malignant B cells preferentially form long 3′UTR-dependent protein complexes that have tumor-promoting roles ^12^. So far, most studies that investigated the functional consequences of alternative 3′UTR isoform expression have relied on using expression constructs, but more recently CRISPR-mediated deletions of 3′UTRs were added to the tool kit ^11-15,25,84-86^. Alternatively, as we showed here, 3′UTR isoform expression can be altered through enhancer deletion (Fig. 1). This strategy will allow researchers to study the resulting functional consequences at endogenous gene loci and has the advantage of keeping 3′UTR *cis*-elements intact while only changing the relative expression of 3′UTR isoforms. 3′UTR length has expanded substantially during evolution of more complex animals and it correlates with the number of cell types observed in an organism ^87,88^. At the same time, the number of enhancers has increased with organismal complexity ^1^. Therefore, we speculate that increased regulation by enhancers has co-evolved with increased regulation by 3′UTRs to integrate intrinsic and extrinsic signals to change gene and mRNA isoform expression ^2^.

## Acknowledgements

We thank all members of the Mayr lab for helpful discussions and Kristian Helin for critical reading of the manuscript. This work was funded by the NIH Director′s Pioneer Award (DP1-GM123454), Damon Runyon-Rachleff Innovation Award, the Pershing Square Sohn Cancer Research Alliance, and the NCI Cancer Center Support Grant (P30 CA008748).

B.K. designed and performed all experiments regarding endogenous PTEN and measured read-through of the reporter, M.M.F. performed all computational analyses, N.P. and J.L. cloned the reporter constructs and performed the reporter assays, J.L. evaluated the knock-down efficiency of the shRNAs, W.M. performed the mRNA stability experiment of the reporter. C.M. conceived and supervised the project, designed and analyzed the reporter experiments, and wrote the manuscript with input from all authors.

The authors declare no competing interests.

## Methods

### Datasets used

To identify the enhancers and promoters used in this study, levels of acetylated H3K27 and transcription factor binding sites in MCF7 cells were visualized using published ChIP-seq data (GSM946850), generated by the Encode project ^39,40^. Binding of MYC to the *PTEN* promoter was assessed by using published ChIP-seq data (GSE33213).

FASTQ files for bulk RNA-seq samples were obtained from the Sequence Read Archive. Mouse definitive erythroblasts: SRR6946157-9 ^64^, SRR8945139-41,44-45 ^66^, mouse hematopoietic stem cells: SRR7946616-7 ^89^, SRR6458998-9000 ^65^.

### Cell culture

For all experiments, the human breast cancer cell line MCF7 was used which was a gift from the laboratory of Robert Weinberg (Whitehead Institute, Cambridge, USA). MCF7 cells were cultured in DMEM supplemented with 10% FCS and 1% Pen-strep (normal condition). For experiments using acidified media, MCF7 cells were cultured for 24 hours in DMEM supplemented with HCl (pH = 6.5). MCF7 cells were transfected with Lipofectamine 2000 (Invitrogen).

#### *PTEN* enhancer deletion using CRISPR-Cas9

All primers are listed in Supplementary Table 3. The guide RNAs targeting the 5′ and 3′ ends of the *PTEN* enhancer were designed using GuideScan and cloned into pX330 as previously described ^90,91^. MCF7 cells were transfected with pmaxGFP and the two pX330 plasmids. After 4 days, GFP-positive cells were sorted into 96-well plates and grown into single cell colonies. Enhancer deletion was tested by PCR and sequencing.

#### shRNA knock-down experiments

shRNAs were designed using the siRNA selection program from the Whitehead Institute and cloned into pSUPERretropuro. Retroviral particles were obtained as described before ^82^. Knock-down efficiency was tested by RT-PCR with gene-specific primers and primers for *GAPDH*.

### Northern blotting

Northern blotting was performed as described previously with modifications ^82^, dx.doi.org/10.17504/protocols.io.bqqymvxw. Briefly, total RNA was isolated using Tri reagent (Invitrogen). PolyA+ mRNA was purified with Oligotex (Qiagen) and 2 µg of polyA+ mRNA was loaded in each lane. The single-stranded probe was generated by unidirectional PCR reaction as described ^92^ with a slight modification. As input for the unidirectional PCR 30 ng of DNA template was used. This consisted of a 724-bp fragment of the *PTEN* coding region amplified from MCF7 cDNA (PTEN-NB-F and PTEN-NB-R). The unidirectional PCR reaction (20 µl) was conducted in buffer (100 mM Tris-HCl/pH 8.3, 500 mM KCl, 15 mM MgCl_2_, 1% Triton-X-100) containing 0.2 mM each of dCTP, dGTP, and dTTP, 0.5 unit of Taq polymerase, the reverse primer from above, and 6 µl of 3000 Ci/mmol [α-^32^P]dATP (Perkin Elmer). The reaction mixture was initially boiled for 10 min at 95°C and subjected to 35 thermal cycles (95°C for 30 s; 45°C for 30 s; 72°C for 1 min), which was followed by 5 min incubation at 72°C. After the PCR reaction, 5 µl of 0.2 mM EDTA was added to the mixture and boiled for 5 min at 95°C, followed by 2 min chilling on ice. The PCR product was then used for hybridization to probe the *PTEN* transcript.

#### mRNA stability of endogenous *PTEN* 3′UTR isoforms

MCF7 WT and dE2 cells were treated with or without 25 mM HCl overnight. The cells were then treated with either DMSO or 4 μg/ml actinomycin D for the indicated time points. Northern blots were performed and the levels of *PTEN-SU* and *PTEN-LU* isoforms were quantified using Multi Gauge (Fuji) and normalized to the levels of *GAPDH* mRNA.

### RT-PCR and PCR

#### *PTEN* enhancer deletion

Genomic DNA was isolated from each clone and the deletion of the *PTEN* enhancer was assessed by PCR with primers (PcE-FS2 and PcE-RS2) flanking the enhancer region and Sanger sequencing. As MCF7 cells are not diploid at the *PTEN* locus, the presence of the alleles with *PTEN* enhancer sequences was further assessed by qPCR using genomic DNA isolated from WT and dE cells to amplify two regions outside of the *PTEN* enhancer (chr10:89,618,718-89,618,843 and chr10:89,620,263-89,620,387) and two regions inside *PTEN* enhancer (chr10:89,621,562-89,621,699 and chr10:89,621,320-89,621,440) using FastStart universal SYBR green master mix (Roche). The qPCR results from the amplicons inside the *PTEN* enhancer were normalized to those from the amplicons outside of the *PTEN* enhancer and were compared between WT and enhancer deletion cells to determine the extent of enhancer loss.

#### Steady-state *PTEN* mRNA

Total RNA was isolated using Tri reagent (Invitrogen) followed by cDNA synthesis using qScript cDNA supermix (Quantabio). To quantify steady-state *PTEN* mRNA level, primers PTEN-qP-F and PTEN-qP-R were used, and qRT-PCR was performed using FastStart universal SYBR green master mix (Roche). *RPL19* mRNA was used as loading control.

#### *PTEN* splicing pattern

A primer pair (PTEN-sp-F and PTEN-sp-R) located in the first and last exon of *PTEN* (NM_000314) was used on the cDNA described above and visualized on 1% agarose gels.

#### Transcript production rate for endogenous *PTEN*

The nascent transcripts were extracted following the protocol described by Russo et al. (2017) ^93^ with minor modifications. Briefly, MCF7 cells were treated with 400 µM 4-thiouridine for 2 hours (4sU, Fisher Scientific). After labeling, Tri reagent (Invitrogen) was used to extract total RNA from cells, followed by DNase I treatment (NEB). Next, 50 µg of total RNA was mixed with 0.5 µg of 4sU-labeled yeast RNA as a spike-in. The mixture was biotinylated with 10 µg of MTSEA biotin-XX (Biotium) in 100 mM HEPES/pH 7.5 and 10 mM EDTA for 30 min at 25°C in dark with gentle agitation and the biotinylated RNA was recovered by ethanol precipitation. Then, half of the biotinylated RNA was mixed with 100 µM DTT and retained as the total RNA fraction. The other half was incubated with Dynabeads M-280 Streptavidin (Invitrogen) in bead wash buffer (100 mM Tris-HCl/pH 7.5, 1 M NaCl, 0.1% Tween-20, 10 mM EDTA), washed twice and eluted twice in 100 µl of 0.1 M DTT at 25°C for 5 min, followed by ethanol precipitation. cDNA generated using Superscript III reverse transcriptase (Invitrogen) and random hexamers (Invitrogen). *PTEN* transcript abundance was measured by qPCR using an amplicon in the first intron of the *PTEN* gene (PTEN-int1-qP-F and PTEN-int1-qP-R) normalized to the abundance of yeast *ACT1* transcripts from the spike-in RNA (using primers yACT1-qP-F and yACT1-qP-R). The rate of transcript production was determined by the ratio of the abundance of the nascent *PTEN* transcripts to the total *PTEN* transcripts.

#### Measurement of read-through transcripts of the reporter constructs

Luciferase reporter constructs were linearized by BglII, which cuts upstream of the enhancer/promoter in the renilla luciferase constructs. 650 fmol of the linearized plasmids were transfected as described above. Total RNA was extracted after 24 hours using Tri reagent (Invitrogen), followed by DNase I (NEB) treatment and ethanol precipitation. 1 µg of total RNA was used to generate cDNA using Superscript III reverse transcriptase (Invitrogen) and random hexamers (Invitrogen). Read-through transcripts were detected with a primer pair that localizes to the vector backbone (1040 bp downstream of the PAS; RenRT-qP-F and RenRT-qP-R). Read-through values were normalized to total transcript levels obtained by a primer pair that localizes to the renilla open reading frame (941 bp upstream of the PAS; RenORF-qP-F and RenORF-qP-R).

#### mRNA stability of the reporter constructs

Luciferase plasmids were transfected as described above. After 24 hours, cells were either treated with DMSO or with actinomycin D (4 µg/ml; Sigma) for the indicated time points. Total RNA was extracted using Tri reagent and was used to generate cDNA using qScript qScript cDNA supermix (Quantabio). qRT-PCR was performed using FastStart universal SYBR green master mix (Roche) on a 7500 HT Fast Real-Time PCR System (Applied Biosystems). The primer pairs used to quantify total *PTEN* mRNA (PTEN-stability-F and -R), luciferase mRNA (Rluc-stability-F and -R) and *GAPDH* mRNA (GAPDH-F and GAPDH-stability-R) (for normalization) are listed in Supplementary Table 3.

#### 5’ RACE

Renilla luciferase plasmids were transfected as described above and total RNA was extracted after 24 hours using Tri reagent (Invitrogen). 5’ RACE was performed with the 5’ RACE kit (Roche) using gene-specific reverse primers: PTEN-5’RACE-R and R2 (for nested PCR).

### Western blotting

Cell pellets were lysed in 2x Laemmli buffer (Alfa Aesar) and performed as described previously ^12^ with the following antibodies: anti-PTEN (A2B1, Santa Cruz Biotechnology, sc-7974, 1:1000) and anti-GAPDH (V-18, Santa Cruz Biotechnology, sc-20357, 1:500). As secondary antibodies anti-mouse IRDye 800 (1:5000; Li-Cor Biosciences, Cat# 926-68072) and anti-goat IRDye 680 (1:5000; Li-Cor Biosciences, Cat# 926-32224) were used.

### Luciferase reporter assays

#### Luciferase reporter constructs

All luciferase constructs were derived from PIS1 vector ^82^. It contains the thymidine kinase promoter of Herpes simplex virus, followed by a renilla luciferase open reading frame, followed by the late SV40 PAS. To obtain reporter constructs with different promoters or PAS, the promoter or the SV40 PAS were exchanged using restriction enzyme digest or Gibson cloning. To obtain reporter constructs with different enhancers, the enhancer sequences (Supplementary Table 1) were cloned upstream in the sense orientation of the respective promoters. In the case of Penh1, the reverse complement of the sequence was cloned downstream of the PAS. MYC-binding sites (E-boxes with the sequence CACGTG at positions -1149 bp and -1353 bp upstream of the transcription start site of the *PTEN* gene) were mutated to CAAGAA using Quikchange Lightening kit (Agilent). The TATA synthetic promoter sequence was derived from pGL firefly reporter with a minimal promoter.

#### Luciferase assay

Luciferase assays were performed in 24-well plates as described previously ^82^. The number of experiments listed in the figures corresponds to biological replicates. In each well, 100 ng of firefly luciferase control reporter plasmid PISO together with 400 ng of renilla luciferase plasmid were transfected. Same molar amounts of plasmid were transfected to account for different construct sizes (400 ng were used for a plasmid of 5000 bp). Firefly and renilla luciferase activities were measured with the Dual-luciferase assay (Promega) 24 hours after transfection. Renilla activity was normalized to firefly activity to control for transfection efficiency. Transcriptional activity of a promoter corresponds to renilla luciferase activity (normalized by firefly luciferase activity) after transfection of the reporter containing the promoter and the SV40 PAS. CPA activity of a test PAS was obtained by dividing the luciferase activities of the constructs with the test PAS by the SV40 PAS in the context of the same promoter.

To assess CPA activity after knock-down of transcription factors, luciferase constructs were transfected into MCF7 cells stably expressing control (ctrl) shRNAs or shRNAs targeting specific transcription factors or co-activators. PAS usage was calculated as described above. When several shRNAs against a specific factor were available, the results were pooled.

### Association of cell type-specific enhancers with alternative 3′UTR isoforms

#### Dataset selection

To obtain genes regulated by cell type-specific enhancers, we used a publicly available dataset generated by Cai et al. (2020) who determined cell type-specific enhancers in murine definitive erythroblasts (here: Ery) and assigned them to target genes using a combination of methods including transcriptome analysis, chromatin accessibility, histone modifications, transcription factor occupancy, and 3D chromatin interactions ^64^. We used the assigned target genes from Supporting Information Dataset_S01.

To identify genes that increase their expression in erythroblasts, we compared gene expression between erythroblasts and murine hematopoietic stem cells (HSC). We set out to use replicates (two HSC datasets ^65,89^ and two Ery datasets ^64,66^). The accession numbers are listed above. To perform quality control for the samples, we aligned them to mm10 using HISAT2 v2.1.0. Gene body coverage was estimated against the mm10 housekeeping gene annotation provided by RSeQC using the geneBody_coverage.py script from RSeQC v4.0.0 ^94^. MultiQC v1.10.1 was used to compare coverages ^95^. This analysis revealed uneven coverage among the samples (Extended Data Fig. 5a). Uneven coverage will confound the analysis on 3′UTR isoform expression as QAPA determines 3′UTR isoform ratios by using the coverage of reads that fall into the region of the *SU* and *LU* isoforms. Moreover, the different datasets use different library preparation methods (pair-end reads, single-end reads, stranded, not stranded) which result in technical differences across the samples independent of their cell type-specific differences. To identify technically similar datasets, we performed principal component analysis (using the top 1000 genes of highest variance in the gene expression matrix after DESeq2’s rlog transformation) on the four datasets. We observed that the datasets generated by Ling (2019) and Lee (2018) had a minimal distance in PC2 which reflects technical variation (Extended Data Fig. 5b) ^65,66^. Therefore, these samples were used for downstream pairwise analysis as representative Ery and HSC cell types, respectively.

#### Differential gene and 3′UTR isoform analysis

To identify genes with a significant difference in gene expression between Ery and HSC, we used DESeq2 v1.28.0 ^96,97^. To identify genes with a significant difference in 3′UTR isoform expression, we first used QAPA to identify multi-UTR genes and to obtain TPM values for *SU* and *LU* isoforms ^98^. Samples were pseudoaligned for transcript quantification in Salmon v1.3.0 (‘salmon quant --gcBias --validateMappings -l A‘) to the pre-compiled QAPA v1.3.0 mm10 3’ UTR annotation with the full mm10 genome as decoy ^99^. Multi-UTR genes used in the analysis were filtered based on Num_Events > 1 (more than one annotated isoform) and at least 3 TPM in one or more samples. Single-UTR genes have Num_Events = 1. *SU* isoforms were identified by QAPA APA_IDs ending in “_P”. To identify statistically significant changes in 3′UTR isoform usage we used DEXSeq v1.34.0 with 10% FDR. Normalized expression values were computed by rescaling QAPA TPM values per sample by estimated size factors from DESeq2. We further required a minimal fold change >2 for *SU* isoform expression to be considered significantly upregulated. All analysis was performed in R v4.0.2 and Bioconductor v1.13.

#### Multi-UTR gene categories

The four multi-UTR gene categories were determined as follows. ‘Gene up’: DESeq2 results (10% FDR, at least 2-fold upregulated in Ery) and no significant 3′UTR ratio change. ‘SU up’: Significant 3′UTR ratio change with at least 2-fold upregulation of *SU* isoform expression using QAPA TPM values together with DEXSeq adjusted p-values and no significant upregulation of gene expression (using DEseq2). ‘SU & gene up’: Genes have a significant 3′UTR ratio change with at least 2-fold upregulation of the *SU* isoform and they have a significant upregulation (at least 2-fold) in gene expression measured by DEseq2. ‘Rest’: is rest of the expressed (TPM >3) multi-UTR genes. Only a few genes showed an exclusive change in *LU* isoform expression (*N* = 28) and were not classified into their own group. Our code will be available at: https://github.com/mfansler/utr-enhancers-pipeline

### Statistics

For all pairwise comparisons of PAS usage or transcriptional activity a 2-tailed, 2-sample unequal variance t-test for independent samples was applied. When comparing several samples, a One-way ANOVA was performed. Chi-square (X^2^) tests were used to test for significant enrichment of genes associated with Ery-specific enhancers. The Pearson value was reported.

## Data availability

The enhancer deletion and control clones will be available upon request. Our code will be available at https://github.com/mfansler/utr-enhancers-pipeline.

## Extended Data Figure legends

**Extended Data Fig. 1:**
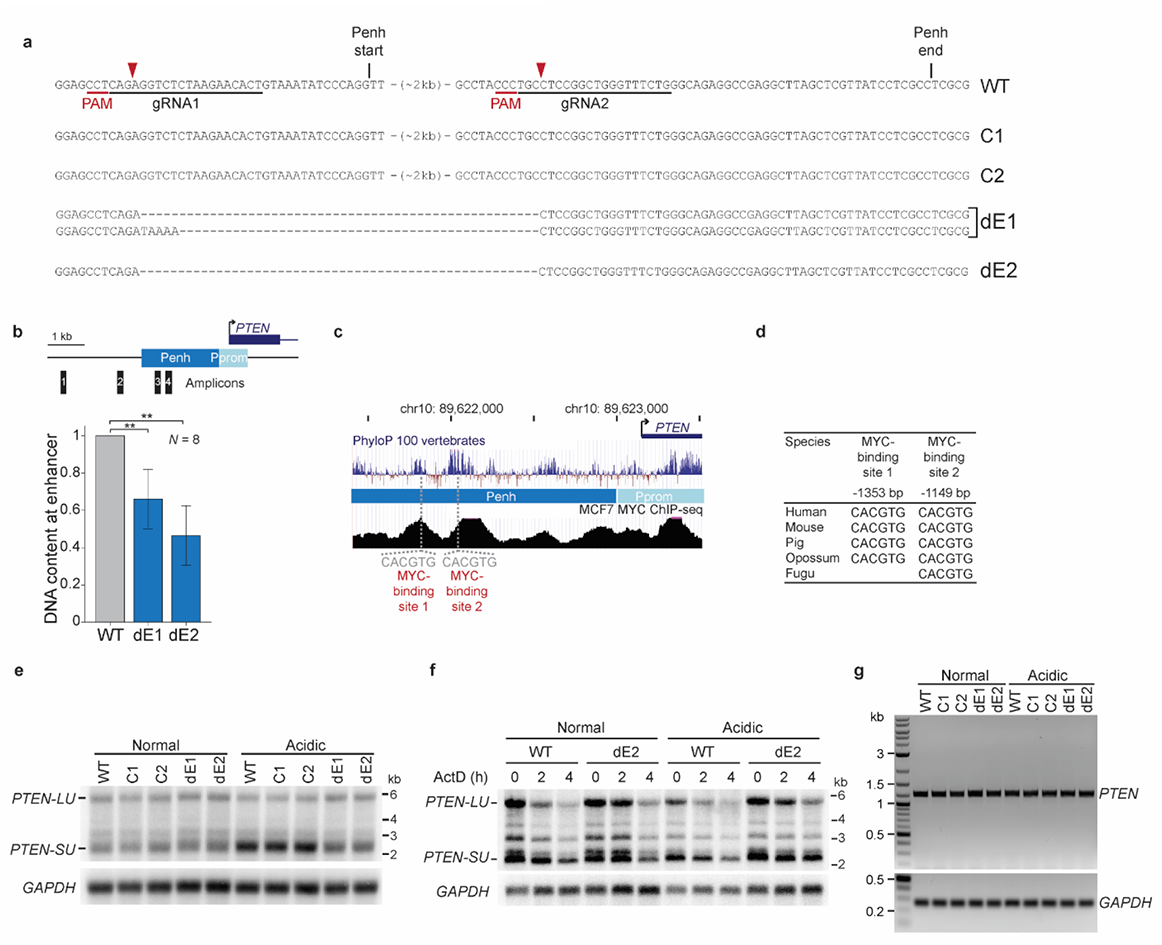
CRISPR-mediated heterozygous deletion of the *PTEN* enhancer in MCF7 cells. **(a)** Sequence alignment of *PTEN* alleles spanning the enhancer deletion sites in parental WT, control clones (C1, C2), dE1 and dE2 clones analyzed in this study. Binding sites of gRNAs used to generate the deletion are underlined in the WT reference sequence and predicted cutting sites are marked by red arrow heads. **(b)** Schematic showing strategy to assess the number of deleted alleles upon CRISPR-mediated *PTEN* enhancer deletion. As MCF7 cells contain more than two alleles for chromosome 10, four PCR amplicons were designed and the DNA content within the deletion was compared to the outside region and normalized to the WT sample. Shown is the fold change in DNA content in the *PTEN* enhancer region in dE1 and dE2 compared with WT cells. One-way ANOVA, *P* = 3 × 10^−7^, t-test for independent samples: WT vs dE1: *P* = 0.001, WT vs dE2: *P* = 3 × 10^−5^. **(c)** UCSC genome browser snapshot showing the *PTEN* gene locus with ChIP-seq data for MYC and the sequence conservation track of 100 vertebrates. The position of the two conserved MYC-binding sites (canonical E-boxes) are indicated. **(d)** Sequence conservation of the two MYC-binding sites located in the *PTEN* enhancer in different organisms. **(e)** Representative northern blot of *PTEN* transcripts from WT, C1, C2, dE1, and dE2 cultivated in normal or acidic conditions. *GAPDH* serves as loading control. **(f)** Northern blot to determine the stability of *PTEN-SU* and *PTEN-LU* transcripts in WT and dE2 cells cultivated in normal or acidic conditions after actinomycin D (ActD) treatment for the indicated time points. *GAPDH* serves as the loading control. **(g)** RT-PCR for full-length *PTEN* mRNA in the indicated samples. The *PTEN* mRNA from the start to the stop codon was amplified. *GAPDH* serves as loading control.

**Extended Data Fig. 2:**
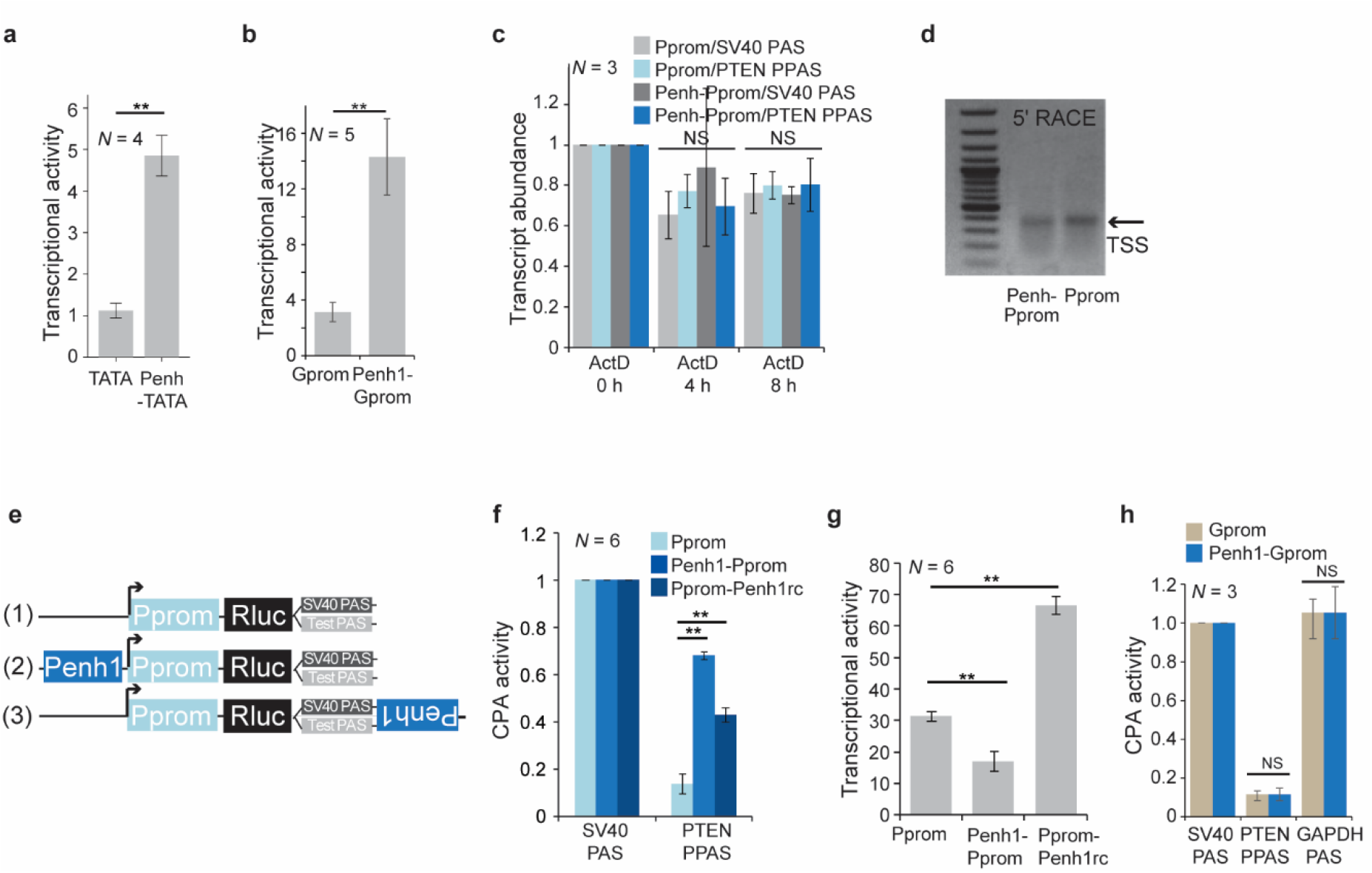
Enhancer-mediated regulation of CPA activity. **(a)** Transcriptional activity of a minimal synthetic promoter (TATA) was measured in the absence or presence of the *PTEN* enhancer (Penh-TATA). As in Fig. 2b. T-test for independent samples was applied. **, *P* = 2 × 10^−4^. **(b)** Transcriptional activity of the *GAPDH* promoter (Gprom) in the presence or absence of the Penh1. As in Fig. 2b. **, *P* = 5 × 10^−4^. **(c)** Reporter mRNA transcript stability was assessed after actinomycin D (ActD) treatment of MCF7 cells expressing the indicated reporters at the indicated time points. The Penh did not influence mRNA stability of the reporters. The sequence context of the SV40 PAS and the PTEN PPAS did not influence the stability of the reporters. Shown is mean ± std. One-way ANOVA was performed. **(d)** 5′RACE was used to determine the transcription start sites of the Pprom reporters in the presence or absence of the *PTEN* enhancer. The canonical transcription start site (TSS) is used in both reporters and is indicated by the arrow. **(e)** Schematic of luciferase reporter constructs to investigate position-dependent enhancer-mediated CPA cleavage activity. The reverse complement (rc) of Penh1 was cloned downstream of the PAS. Shown as in Fig. 2a. **(f)** PTEN PPAS cleavage activity measured using the reporter constructs shown in (e). Shown is mean ± std. T-test for independent samples was performed; **, *P* = 1 × 10^−5^. **(g)** Transcriptional activity of the reporters from (e) shown as in Fig. 2b. T-test for independent samples was performed; **, *P* = 1 × 10^−5^. **(h)** Enhancer-dependent CPA activity in the context of the Gprom in the presence or absence of the Penh. As in Fig. 2c.

**Extended Data Fig. 3:**
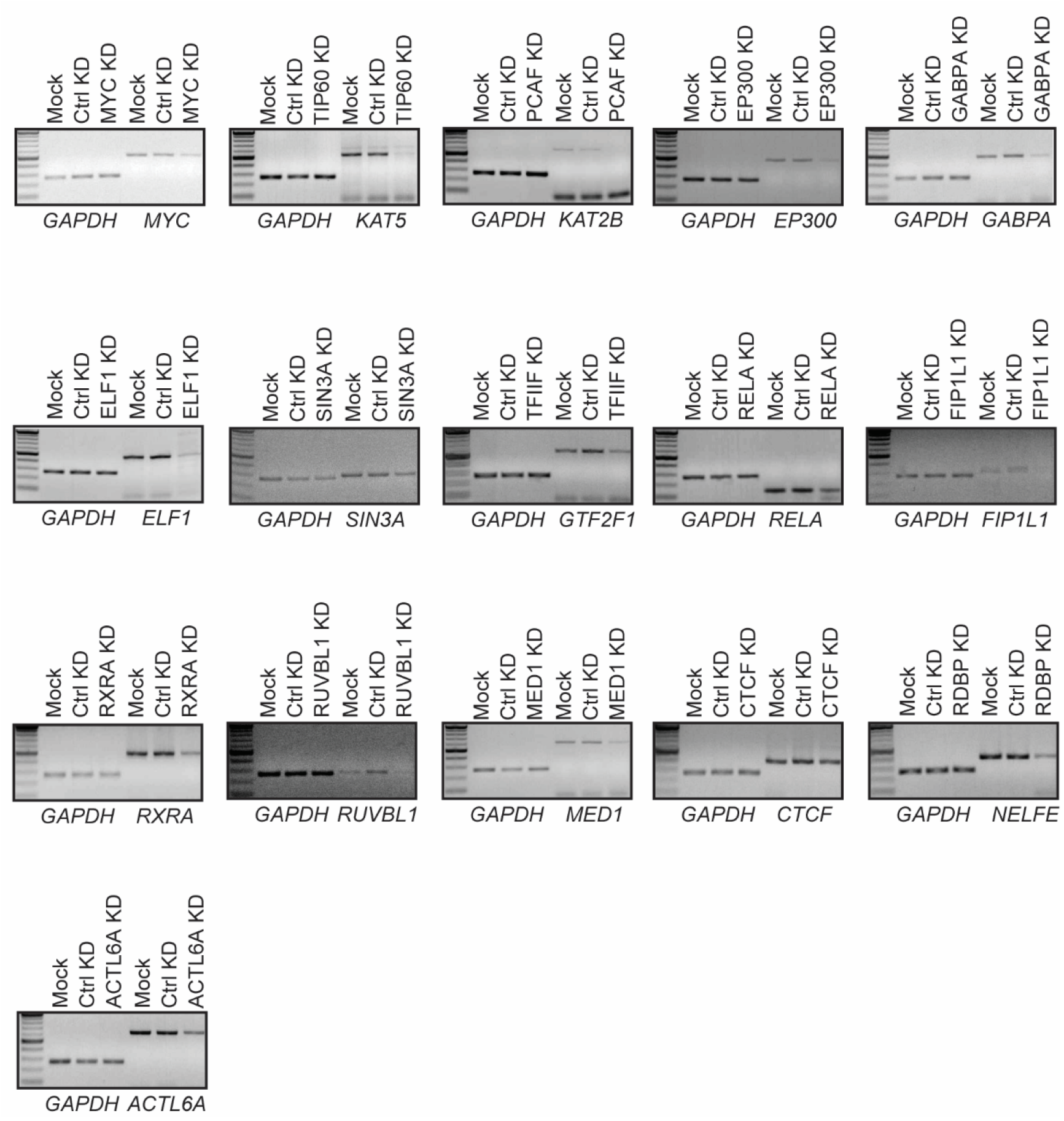
shRNA-mediated knock-down of transcription factors and co-activators in MCF7 cells. Shown are mRNA levels of transcription factors and co-activators in MCF7 cells, after stable expression of a control shRNA (Ctrl KD) or an shRNA against the indicated factor. Mock, no shRNA was transfected. *GAPDH* serves as loading control.

**Extended Data Fig. 4:**
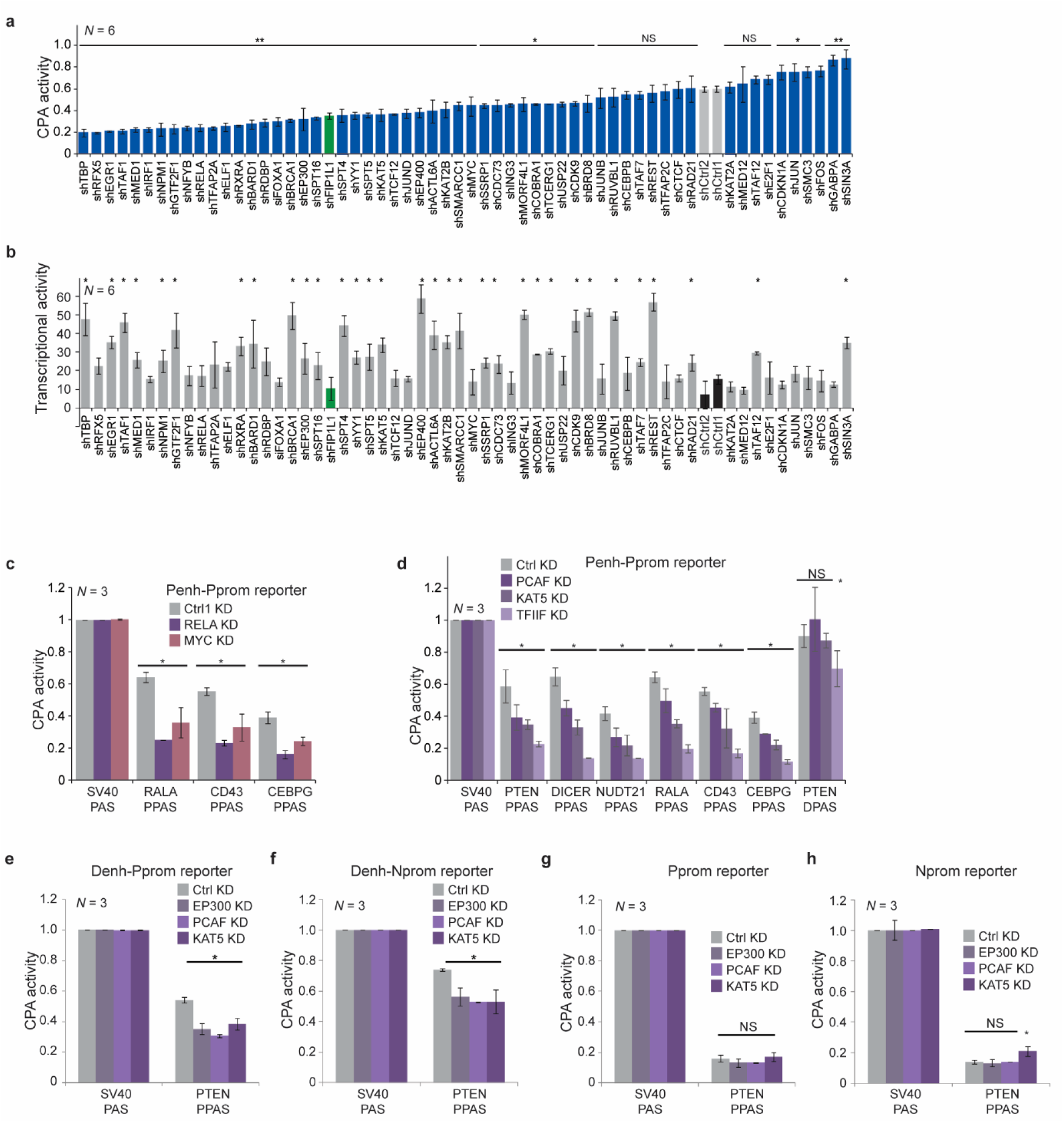
shRNA screen identifies transcription factors and co-activators that regulate CPA activity of the PTEN PPAS. **(a)** CPA activity of the PTEN PPAS in the context of the Penh-Pprom reporter after KD of the indicated transcription factors or co-activators is shown as mean ± std. KD of FIP1L1 (green; polyadenylation factor) serves as positive control. CPA activity in ctrl KD is similar to CPA activity of the Penh-Pprom reporter in Fig. 2c. T-test for independent samples was performed; **, *P* < 0.001; *, *P* < 0.02. See Supplementary Table 2 for values. **(b)** As in (a) but shown is transcriptional activity. *, indicates a change in transcriptional activity > 2-fold, compared to the average of the controls (ctrl1 and ctrl2). **(c)** CPA activity of additional PAS after KD of transcription factors (as in Fig. 4d). T-test for independent samples was performed; *, *P* < 0.01. **(d)** CPA activity of additional PAS after KD of co-activators (as in Fig. 4d). T-test for independent samples was performed; *, *P* < 0.02. **(e-h)** CPA activity of the PTEN PPAS in the context of different promoters in the presence or absence of the distal enhancer after of KD of histone acetyl transferases. Shown is mean ± std. T-test for independent samples was performed. **, *P* < 0.001; *, *P* < 0.03.

**Extended Data Fig. 5:**
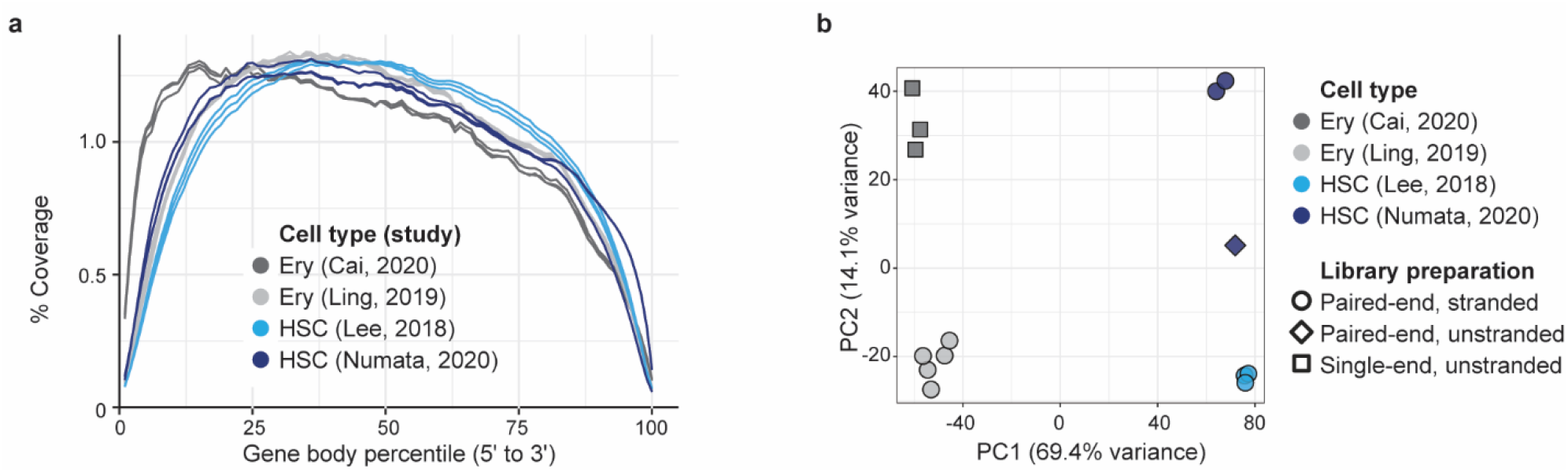
Quality controls for the used RNA-seq datasets. **(a)** RNA-seq gene body coverage of four datasets on murine erythroblasts (Ery) and hematopoietic stem cells (HSC). **(b)** Principal component analysis on the datasets from (a). In addition to variation caused by cell type-specific differences in gene expression (PC1), two of the datasets show large technical variations likely caused by different library preparation methods (PC2). The datasets with the least difference in technical variation (Lee, 2018 and Ling, 2019) were used for the analysis in Fig. 5.

---

**Supplementary Table 1.**
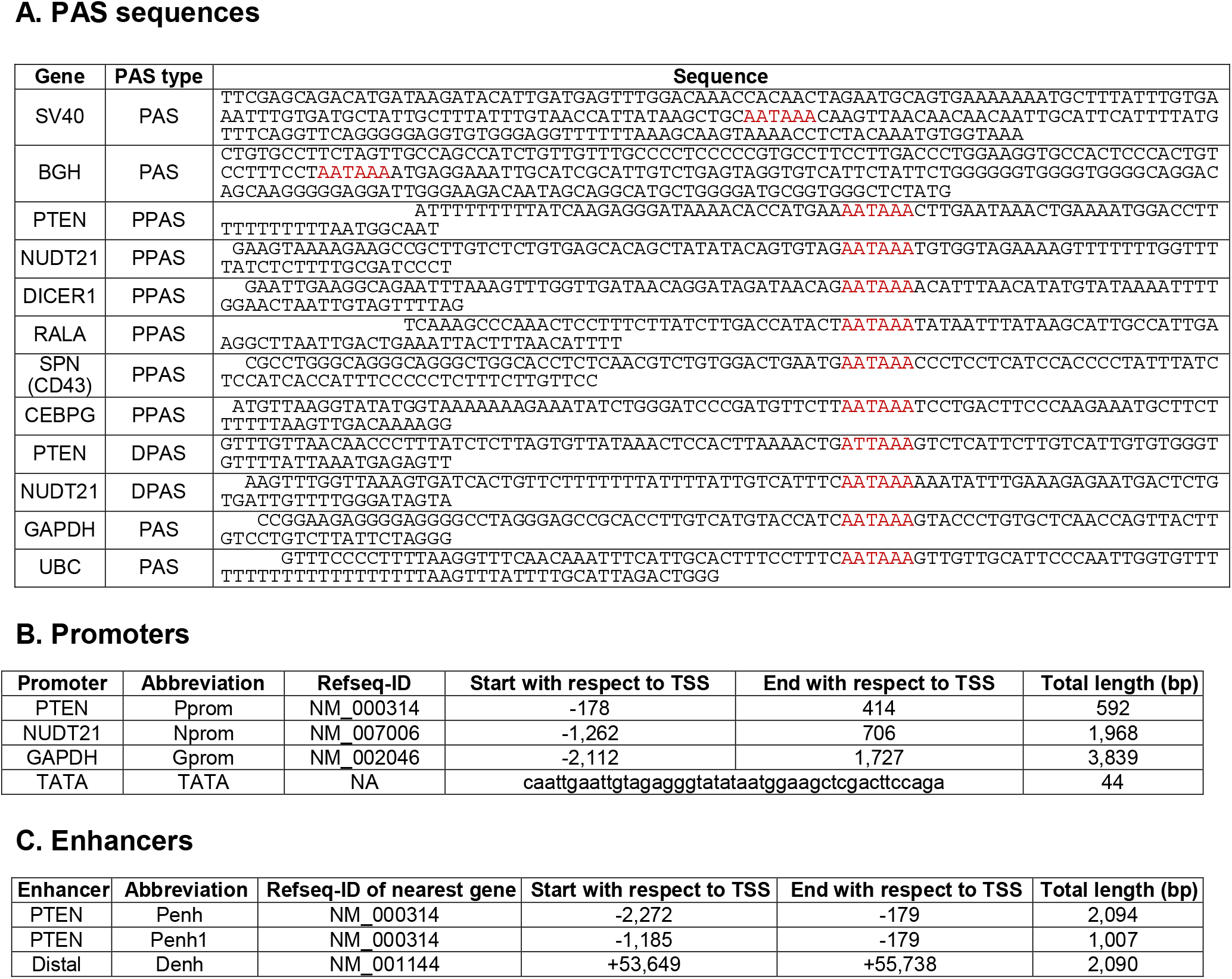
Sequences of PAS, promoters, and enhancers.

**Supplementary Table 2.**
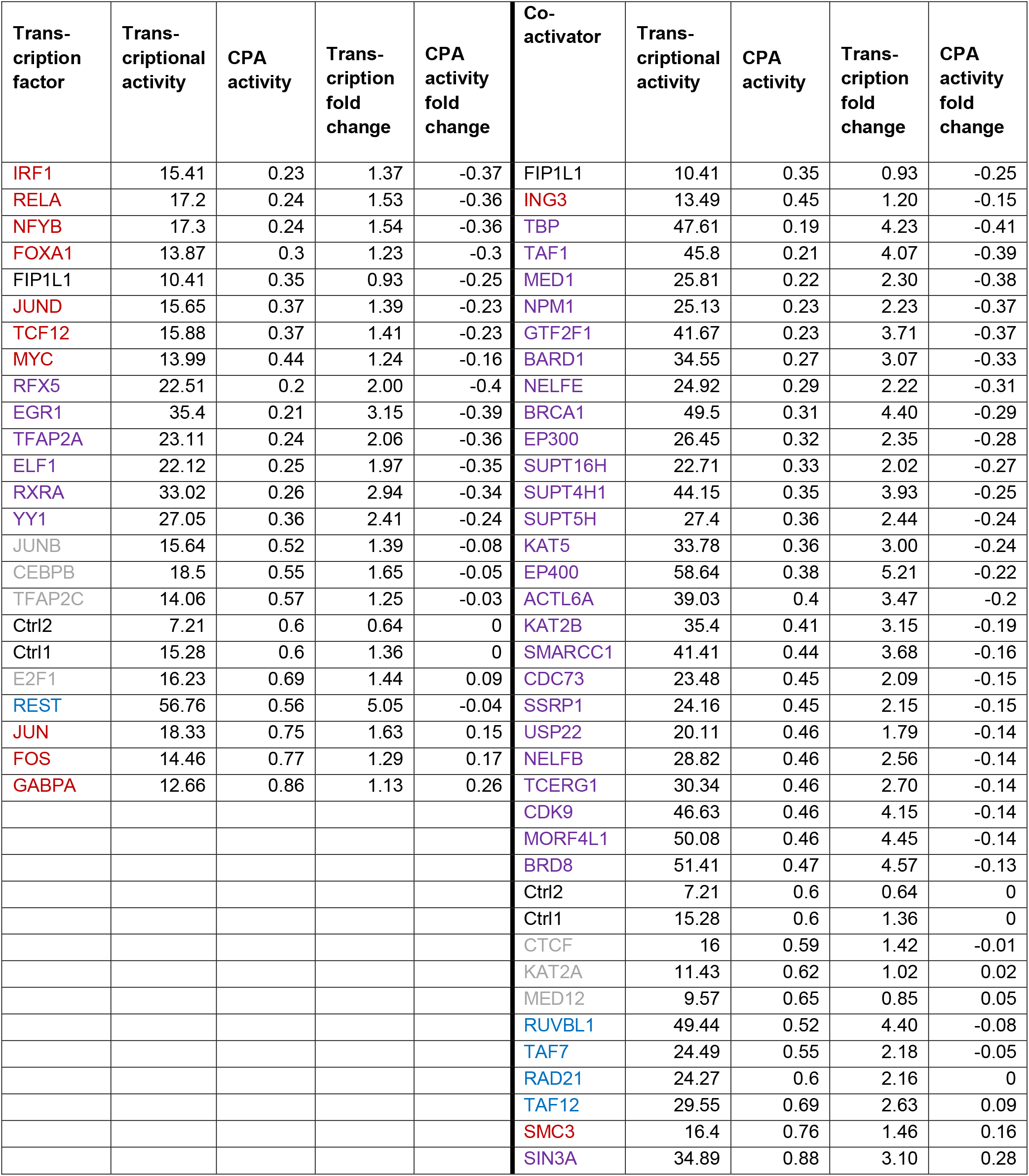
Results of the shRNA screen.

**Supplementary Table 3.**
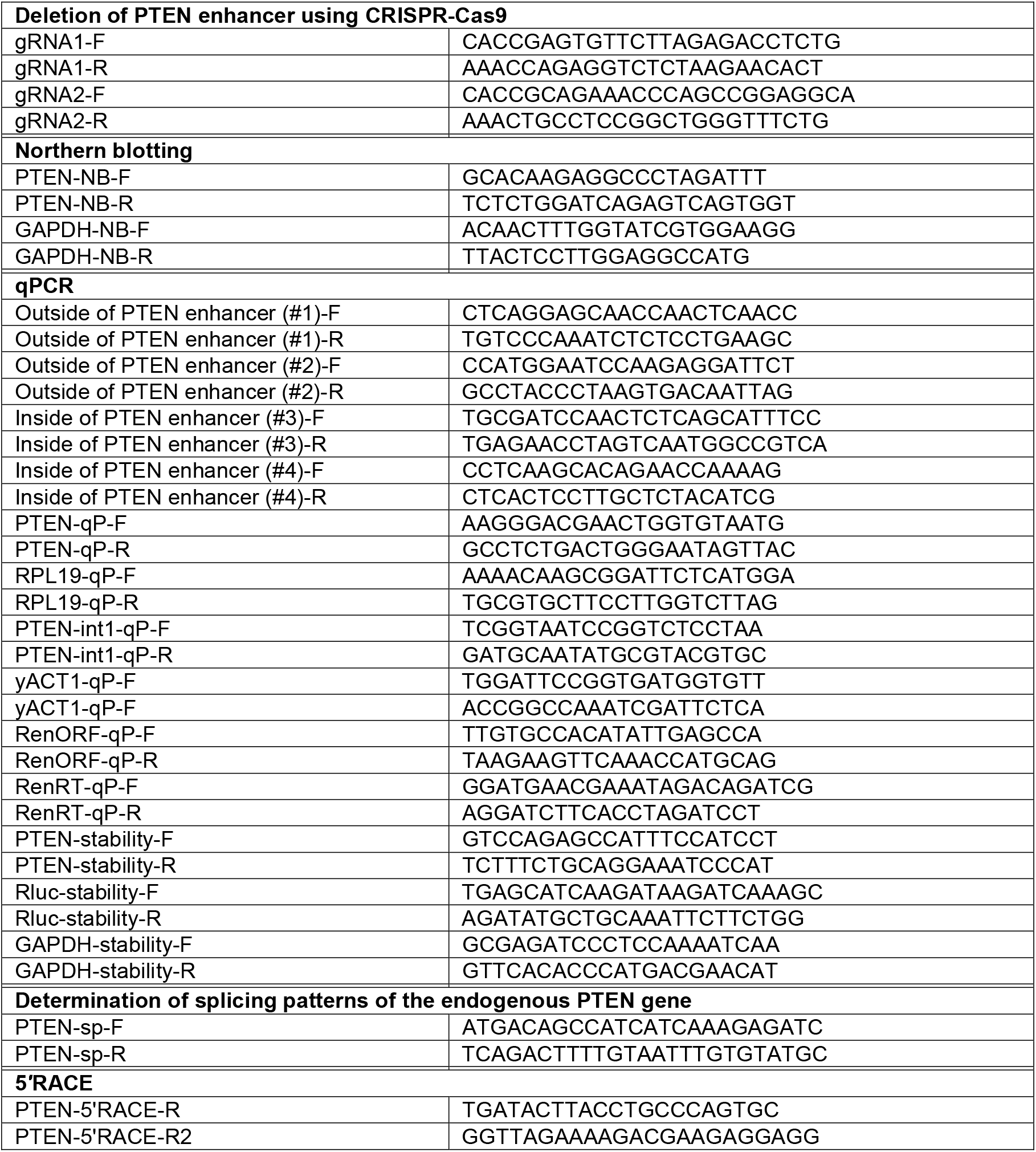
Primer sequences.

